# Genomic analysis of extended-spectrum beta-lactamase (ESBL) producing *Escherichia coli* colonising adults in Blantyre, Malawi reveals previously undescribed diversity

**DOI:** 10.1101/2021.10.07.463523

**Authors:** Joseph M. Lewis, Madalitso Mphasa, Rachel Banda, Mathew A. Beale, Jane Mallewa, Catherine Anscome, Allan Zuza, Adam P. Roberts, Eva Heinz, Nicholas R. Thomson, Nicholas A Feasey

## Abstract

*Escherichia coli* is a ubiquitous bacterium and one of the most prevalent Gram-negative species associated with drug resistant infections. The large number of sequenced genomes available have provided us with a consistently growing knowledge base to further understand pathogenesis and epidemiology of this organism. However, data from sub-Saharan Africa (sSA) are underrepresented in global sequencing efforts and *E. coli* genetic diversity from this region is poorly described. To reduce this gap, we investigated extended-spectrum beta-lactamase (ESBL)-producing *E. coli* colonising adults in Blantyre, Malawi to assess the bacterial diversity and AMR determinants and to place these isolates in the context of the wider population structure. We performed short-read whole-genome sequencing of 473 colonising ESBL *E. coli* isolated from human stool and contextualised the genomes with a previously curated multi-country species wide collection of 10,146 genomes. The most frequently identified sequence types (STs) in our collection were the globally successful ST131, ST410 and ST167, and the dominant ESBL genes were *bla*_CTX-M_, mirroring global trends. However, 37% of Malawian isolates did not cluster with any isolates in the curated multicountry collection and a core gene phylogeny was consistent with locally spreading subclades within globally dominant clones, including in ST410 and ST167. We also found Carbapenemase genes in our collection at low frequency; we used long read sequencing to characterise selected ESBL and carbapenemase-associated plasmids, demonstrating the presence of globally distributed carbapenemase carrying plasmids. Increased genomic surveillance of *E. coli* from Malawi and sSA is necessary to understand local, regional and global transmission of both *E. coli* and the AMR genes they commonly carry.

**Impact Statement:** Drug-resistant *Escherichia coli* producing extended-spectrum beta lactamase (ESBL) or carbapenemase enzymes have been identified by the World Health Organisation as priority pathogens of global concern, and whole genome sequencing has provided insight into mechanisms of virulence, antimicrobial resistance, and the spread of high-risk clones. However, studies analysing large numbers of *E. coli* using whole-genome data often focus on opportunistic use of hospital diagnostic collections in high-income settings. Understanding how the genomic epidemiology of *E. coli* in low- and middle-income countries (including many of the nations of sub-Saharan Africa) differs is essential to provide insight into local, and global drivers of transmission. We therefore sequenced 473 ESBL-producing *E. coli* genomes colonising adults in Blantyre, Malawi. We analyse determinants of antimicrobial resistance and virulence and place the isolates in wider context using a previously published global *E. coli* collection that was generated to represent the whole species diversity of sequences publicly available at the time of generation. We find that there is diversity in Malawian isolates not reflected in the curated global collection: widely successful antimicrobial-resistance associated *E. coli* sequence types are represented in Blantyre, but locally circulating subclades are apparent. Furthermore, given the high number of ESBL producing pathogens causing infections there is an unmet need for carbapenem antimicrobials which are still active against ESBL-producers but are not yet widely available in our setting. We find that carbapenemases (enzymes that can render bacteria resistant to carbapenems) in our collection are unusual but present and carried on globally disseminated plasmids. So too are globally successful, stably carbapenemase-associated *E. coli* lineages. Although the Malawian isolates analysed typically lacked carbapenemases, carbapenem use is increasing in Malawi and their unstewarded use will accelerate selection for carbapememases in *E. coli* in the future. Our study highlights the need for robust stewardship protocols and ongoing genomic surveillance as these agents are introduced.

**Data Summary:** All data and code to replicate this analysis are available as the *blantyreESBL* v1.3 R package (https://doi.org/10.5281/zenodo.5554081) available at https://github.com/joelewis101/blantyreESBL. Reads from all isolates sequenced as part of this study have been deposited in the European Nucleotide Archive, under PRJEB26677, PRJEB28522 and PRJEB36486 (short reads) and PRJNA869071 (Nanopore reads and hybrid assemblies). Accession numbers (as well as accession numbers of publicly available genomes used in this analysis) linked to sample metadata are provided in the R package and as supplementary data to this manuscript.

## Introduction

*Escherichia coli* is a ubiquitous bacterium, a human gut commensal and common pathogen^1^. Beta-lactam antibiotics (including third-generation cephalosporins, 3GC) are widely used for treatment of Gram-negative infections like *E. coli* but are largely rendered ineffective by bacteria expressing extended-spectrum beta lactamase (ESBL) enzymes. ESBL-producing bacteria have disseminated globally, in many cases leaving carbapenems as the only clinically effective and well tolerated treatment option^2,3^. These agents are now also under threat given the increasing spread of strains producing carbapenem-inactivating carbapenemase enzymes, and *E. coli* producing ESBL and carbapenemase enzymes have been identified as priority pathogens by the World Health Organisation^4^. Global genomic surveillance has provided insight into the mechanisms and epidemiology of their spread, suggesting that capture of virulence and AMR determinants via horizontal gene transfer by so-called high risk clones results in fitness and/or colonisation advantages and subsequent global dissemination^5^. This phenomenon is well described in *E. coli* sequence type (ST) 131^6^, associated with the ESBL-encoding gene *bla*_CTX-M-15_, but has also been recently described in other carbapenemase-associated *E. coli* lineages, such as ST167^7^ and ST410^8^.

Whole genome sequencing efforts to date have largely focussed on AMR *E. coli* collections from high-income settings^9^, however the highest incidence of drug-resistant infection are in low- and-middle income settings, particularly sub-Saharan Africa^10,11^. The genomic epidemiology of AMR *E. coli* in these regions is poorly described. The 3GC antimicrobial ceftriaxone has been widely used in Malawian hospitals since its introduction to the national formulary in 2005^12^, but carbapenem use is not yet routine. Since 2005, ESBL-producing *E. coli* have become an increasing problem in clinical practice and now represent 31% of invasive *E. coli* in Blantyre^13^, whereas carbapenem resistance has so far only been described sporadically.^14^ There is a significant need for better access to carbapenem antimicrobials to treat resistant infections, but the example of ceftriaxone in Malawi and of carbapenems globally shows that carbapenem resistance will likely disseminate rapidly as carbapenem use increases. In this context, both robust antimicrobial stewardship and ongoing genomic surveillance are critical.

In this study, we analysed the genomic diversity of ESBL-producing *E. coli* from a study of gut mucosal colonisation with ESBL Enterobacterales in Blantyre, Malawi, and describe the diversity and AMR determinants of ESBL *E. coli*. We contextualised genomic data from Blantyre with that from large public datasets to understand the diversity of colonising *E. coli* in our setting and to gain insights into how well high-income country (HIC)-focused collections capture and represent the genetic diversity of collections from low- and middle-income-country (LMIC) settings.

## Methods

The clinical study which provided the isolates for this analysis was approved by the Liverpool School of Tropical Medicine Research Ethics Committee (16-062) and University of Malawi College of Medicine Research Ethics Committee (COMREC P.11/16/2063). The isolates analysed in this study were selectively cultured from stool and rectal swabs collected from adults in Blantyre, Malawi, as part of a study of longitudinal carriage of ESBL-producing Enterobacterales, as previously described^15^. Briefly, three groups of adults (≥ 16 years) were recruited: i) 225 adults with sepsis in the emergency department of Queen Elizabeth Central Hospital (QECH), Blantyre, Malawi; ii) 100 antimicrobial-unexposed adult inpatients at QECH; and iii) 100 antimicrobial-unexposed adults recruited from the community. Antimicrobial unexposed was defined as no receipt of antimicrobials in the previous four weeks, with the exception of long-term co-trimoxazole preventative therapy (CPT, trimethoprim-sulfamethoxazole administered lifelong to people living with HIV in Malawi as per World Health Organisation [WHO] guidelines^16^) or antituberculous chemotherapy (usually comprising rifampicin, isoniazid, pyrazinamide and ethambutol). Up to five stool samples (or rectal swab samples) were collected over the course of six months and aerobically cultured overnight at 37ºC on ChromAGAR ESBL-selective chromogenic media (ChromAGAR, France) before being speciated with the API system (Biomeriuex, France).

A subset of isolates identified as *E. coli* underwent DNA extraction and sequencing: one *E. coli* colony pick from the first 507 samples where *E. coli* was identified (this number determined by logistic considerations). DNA was extracted from overnight nutrient broth cultures using the Qiagen DNA mini kit (Qiagen, Germany) as per the manufacturer’s instructions. DNA was sequenced at the Wellcome Sanger Institute on the Illumina HiSeq X10 instrument (Illumina Inc., United States) to produce 150 bp paired end reads. Species was confirmed with Kraken v0.10.6 and Bracken v1.0^17^ with a 8Gb MiniKraken database (3 April 2018). We first reconstructed a core gene phylogeny for the study isolates: *de novo* assembly was performed using SPAdes v3.14.0^18^ and the pipeline described by Page et al^19^, and quality of the assemblies assessed with CheckM v1.1.2^20^ and QUAST v5.0.2^21^. Assemblies with a total assembled length of < 4Mb or with a CheckM-defined contamination of ≥ 10% were excluded from further analysis. Included assemblies had a median 92 (IQR 68-122) contigs and N50 of 180kbp (IQR 123-234kbp). Assemblies were annotated with Prokka v1.5 using a genus-specific database from RefSeq^22^ and the Roary v1.007 pangenome pipeline^23^ used to identify core genes with a BLAST threshold of 95%; paralogs were not split. Genes present in ≥ 99% samples were considered to be core, and a pan-genome of 26,840 genes was determined, of which 2,966 were core. The core genes were concatenated to a 1,388,742 base pseudosequence; 99,693 variable sites were identified and extracted with snp-sites v2.4.1^24^ -c to select for ACGT-only sites and a maximum-likelihood phylogeny inferred from this alignment with IQ-TREE v1.6.3^25^. The ModelFinder module was used to select the best fitting nucleotide substitution model: the general time reversible model with FreeRate site heterogeneity with 5 parameters, which was fitted with 1000 ultrafast bootstrap replicates.

ARIBA v.2.12.1^26^ was used on the reads to identify AMR-associated genes from the SRST2 curated version of the ARG-ANNOT database^27^, and was also used to call single nucleotide polymorphisms (SNPs) in the quinolone-resistance determining regions (QRDR) *gyrA, gyrB, parC* and *parE*, using the wild-type genes from the *Escherichia coli* K-12 substr. MG1655 (NC_000913.3) as reference. Any QRDR mutation conferring quinolone resistance in *E. coli* in the comprehensive antibiotic resistance database^28^ (CARD) was considered to confer quinolone resistance. Beta lactamases were phenotypically classified according to https://ftp.ncbi.nlm.nih.gov/pathogen/betalactamases/Allele.tab. ARIBA was also used to determine *E. coli* multilocus sequence type (ST) as defined by the 7-gene Achtman scheme^29^ hosted at pubMLST (https://pubmlst.org/), to identify plasmid replicons using the PlasmidFinder database^30^, and to determine pathotype by identifying genes contained in the VirulenceFinder database^31^. Pathotype was assigned based on the criteria in Supplementary Table 1^32^. *E. coli* phylogroups were defined by the Clermont scheme using ClermonTyping v20.3^33^.

To place the isolates from this study in context of the wider *E. coli* population structure, we used a dataset from a previously described highly curated multi-country collection of *E. coli* genomes. The collection comprises 10,146 *E. coli* genomes derived from Europe, the Americas, Asia, Africa and Oceana^9^ and was intensely quality-controlled, providing a curated set of genomes representing a species-wide background dataset that we used to bring our samples into context. Henceforth this will be referred to as the global collection. In the original publication describing this collection, the 10,146 genomes were clustered with the popPUNK algorithm into 1,154 lineages and 10 genomes from each of the largest 50 lineages were selected to produce a collection of 500 genomes representative of these largest lineages. Since our Blantyre genomes might not all be members of the 50 largest lineages, to place our isolates in context with this collection we first used popPUNK v1.1.5^34^ to compare our assemblies with the popPUNK database of all 10,146 genomes from the global collection. This allowed us to assign genomes from Malawi into the respective published popPUNK groups. We then constructed a core-gene phylogeny. We combined the 500 curated, representative assemblies of the 50 largest lineages from the global collection with up to 10 genomes from all other PopPUNK-defined lineages in the collection; if a lineage included ≤ 10 genomes we included all of them, otherwise we randomly sampled 10 genomes from the lineage to include. In total, this left a dataset of 2,776 genomes. To include invasive isolates from our setting as comparison to our colonisation isolates, we furthermore added 97 genomes from a previous study of *E. coli* at QECH, where archived samples were selected for sequencing to maximise temporal and antimicrobial susceptibility profile diversity^35^. QC, assembly, determination of ST and phylogroup of these samples proceeded as described above. Following QC, four genomes from this latter study were excluded, and combined with our 473 new Blantyre genomes, this left a final working dataset of 3,342 *E. coli* genomes for analysis. We used the Roary pan-genome pipeline to infer a pangenome, identifying 73,062 gene orthologs in this collection, of which 2,182 were core and formed a concatenated sequence of 52,639 bases with 9,117 variable sites. These were extracted with snp-sites and used to infer the phylogeny, using IQ-TREE with the same substitution model as above, and again with 1000 ultrafast bootstrap replicates.

To better describe the phylogeny of the three dominant STs in our dataset, ST131, ST410 and ST167, in greater resolution, we inferred lineage-specific phylogenies using a map-to-reference approach, contextualised with publicly available genomes. We used previously curated multi-country collections of 862 ST131 genomes^36^, 327 ST410 genomes^8^, and 181 ST167 genomes^37^. For all isolates, we obtained available raw reads (n = 108 genomes for ST167, n = 327 for ST410 and n = 862 for ST131), performed QC with fastQC v0.11.8 (https://www.bioinformatics.babraham.ac.uk/projects/fastqc/) and multiqc v1.8^38^, trimmed raw reads with Trimmomatic v0.39^39^, removing adapter sequences and leading or trailing bases with phred score < 4, bases with a mean score < 20 (over a sliding window of 4 bases), and any reads with length below 36 following removal of low quality bases. We mapped the reads to ST specific references: for ST131 the ST131 NCTC 13441 reference strain from the UK Health Security Agency culture collection (NCBI accession NZ_LT632320.1) and for ST167 and ST410 reference genomes from the curated FDA-ARGOS database^40^ (GenBank accession CP023870.1 for ST167 and CP023870.1 for ST410). We used the snippy v4.6.0^41^ pipeline with default settings, excluding mapped pseudosequences with mean mapped coverage depth <20x from further analysis. Following this additional control; 102 ST167, 326 ST410, and 855 ST131 context genomes were retained for further analyses, and combined with the 38 ST167, 45 ST410 and 64 ST131 from this study. Areas of recombination were predicted with Gubbins v3.0.0^42^ and masked. The variable sites remaining (13,693 sites in a 4,711,093 base alignment for ST410, 6,119 sites in 4,897,877 bases for ST167 and 18,171 sites in 5,174,631 bases for ST131) were used to reconstruct a phylogeny with IQ-TREE as above. Presence of AMR genes and plasmid replicons were inferred as above.

To gain better insights into carbapenemase associated strains in our setting we performed long read sequencing on a *bla*_NDM-5_ carbapenemase associated ST2083 isolate (the only carbapenemase-containing isolate in our collection) and a randomly selected ST410 isolate. This ST was chosen because ST410 are a global disseminated stably *bla*_NDM-5_ associate lineage, but Malawian ST410 isolates (in this study) all lacked the expected carbapenemase genes (see results below). We isolated long-fragment DNA using the epicentre MasterPure™ Complete DNA and RNA Purification Kit as per manufacturer instructions, performed long-read sequencing using a MinION sequencer (Oxford Nanopore) and generated hybrid assemblies. We assembled our genomes using unicycler^43^ v0.4.7, using recommended settings for hybrid assembly of Illumina and nanopore reads. Annotation was performed using prokka as above. To map reads back to the assembly for investigating coverage of AMR regions and ST-specific SNPs in ST410, we used snippy with the default settings. Analysis of the conserved plasmids was performed using BRIG v0.95^44^ with the plasmids CP034954.1, CP034955.1, CP034956.1 and CP034957.1 from the *bla*_*NDM-5*_-encoding ST410 strain SCEC020026^8^ as reference sequences. The unicycler-assembled ST410 strain resulted in nine contigs (4,670,094bp; 127,483bp; 98,229bp; 33,870bp; 2,088bp; 245bp; 194bp; 165bp; 119bp); we did not include the chromosome (largest fragment) and the smallest four plasmids in the comparison as they are likely spurious assemblies; sequences below 200bp furthermore get removed by default by genbank whole-genome upload as likely contaminants. The unicycler-assembled ST2083 strain resulted in five contigs with lengths (4,967,815bp; 110,151bp; 108,044bp; 48,551bp; 46,161bp), all of which were included in the BRIG plasmid comparison with the exception of the chromosome (largest fragment). The assembled ST410 and ST2083 (including all small fragments) and their respective long reads used for the hybrid assembly are deposited under sample accessions SAMN30282368 and SAMN30282369, respectively.

All statistical analyses were carried out in R v4.1.1 (R Foundation for Statistical Computing, Vienna, Austria) and trees were visualized using the *ggtree* v2.2.4^45^ package. Summary statistics, where presented, are medians and interquartile ranges or proportions unless otherwise stated. Short reads from all isolates sequenced in this study have been deposited in the European Nucleotide Archive under project IDs PRJEB26677, PRJEB28522 and PRJEB36486.. All data and code to replicate this analysis are available as the *blantyreESBL* v1.3^46^ R package available at https://joelewis101.github.io/blantyreESBL/. Sample accession numbers linked to sample metadata as well as accession numbers of all context genomes used in this analysis are available as part of the R package and as supplementary data to this manuscript.

## Results

### Population structure

Following quality control, 473 *E. coli* genomes sequenced for this study were included in the analysis, 440 from participants enrolled in hospital, and 33 from community members, with a median 2 (IQR 1-5) samples per participant. A full description of study participants and temporal trends has previously been made^15^. The most common phylogroup was A (43%), followed by phylogroup B2 (20%), B1 (9%), C (9%), F (8%) and D (5%), with nine samples untyped by the Clermont scheme (Figure 1A). The phylogroup distribution differed between isolates from this study and the global collection (Supplementary Figure 1) with a higher proportion of phylogroup A isolates and lower proportion of phylogroup E isolates in our data. Fifty-seven recognised STs were identified in our isolates, with a median of 2 (IQR 1-9, range 1-64) samples per ST (Figure 1B). Examining the core gene tree topology, we confirmed that these STs were largely monophyletic (Supplementary Figure 2). The three most frequent STs accounted for 32% of isolates: ST131 was most commonly identified (64/473 [14%] of isolates) followed by ST410 (45/473 [10%]) and ST167 (38/473 [8%]).

**Figure 1:**
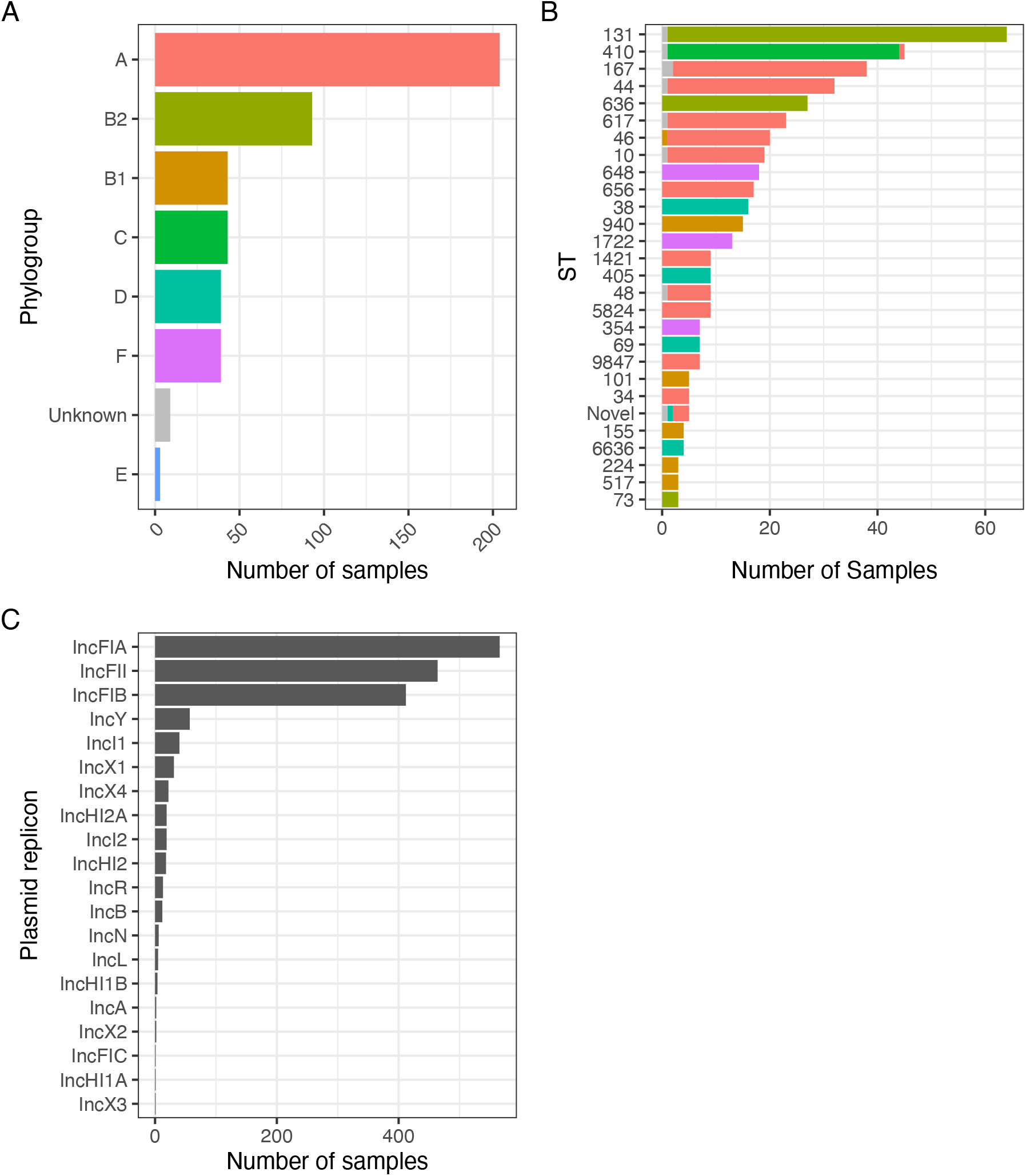
ST (A) and phylogroup (B) distribution of included isolates and (C) identified Inc-type plasmid replicons.

We next placed the carriage isolates from this study in context of the wider species diversity using the global collection^9^. Using popPUNK we assigned our 473 Blantyre isolates to 109/1154 of the clusters (median size 1; IQR 1-3) defined in the original description of the global collection. The distribution of clusters differed between the isolates from this study and the global collection (Figure 2); the largest 50 popPUNK clusters in the global collection contain 76% of all its isolates, but only 140/473 (30%) of isolates from this study. Whilst the largest two popPUNK clusters in this study were commonly represented in the global collection lineage 2 (n = 53 isolates from this study, all ST131) and lineage 40_708 (n = 44 isolates from this study, all ST410), other large clusters in this study had very few representatives in the global collection: the third and fourth largest clusters in our collection were lineage 684 (n = 29 in this study, all phylogroup A ST44) and 451 (n = 27 all phylogroup A ST636) with only one isolate each in the global collection (Table 1). Moreover, a large part of the diversity in our study was not represented in the global collection with 175/473 (37%) of our isolates, forming novel PopPUNK clusters unrelated to isolates in the global collection.

**Table 1:** Phylogroup, sequence type (ST), continent of collection and pathotype of popPUNK-defined clusters

| Cluster | Phylogroup | This study |  | Horesh Collection |  |  |  |
| --- | --- | --- | --- | --- | --- | --- | --- |
|  |  | n | STs | n | STs | Location | Pathotype |
| 2 | B2 | 53 | ST131 (1.00) | 781 | ST131 (0.99) | Europe (0.73);Unknown (0.12);Americas (0.11);Oceania (0.03);Asia (0.00) | ExPEC (0.54);Not determined (0.46) |
| 40_708 | C | 44 | ST410 (1.00) | 29 | ST23(0.41);ST410 (0.34); ST2230 (0.07);ST369 (0.07);ST5491 (0.07);ST5286 (0.03) | Europe (0.62);Americas (0.31);Asia (0.07) | ExPEC (0.55);STEC (0.21);ETEC (0.17);Not determined (0.07) |
| 684 | A | 29 | ST44 (1.00) | 1 | ST44 (1.00) | Europe (1.00) | ExPEC (1.00) |
| 451 | B2 | 27 | ST636 (1.00) | 1 | ST636 (1.00) | Americas (1.00) | ExPEC (1.00) |
| 434_505 | A | 24 | ST167 (0.92);ST10 (0.08) | 3 | ST10 (1.00) | Americas (0.67);Europe (0.33) | ExPEC (0.67);Not determined (0.33) |
| 44_579 | F | 17 | ST648 (1.00) | 26 | ST648 (1.00) | Europe (0.42);Unknown (0.38);Americas (0.19) | ExPEC (0.73);Not determined (0.27) |
| 1186 | A | 17 | ST656 (1.00) | 0 | - | - | - |
| 290 | A | 16 | ST46 (1.00) | 2 | ST46 (1.00) | Americas (1.00) | ExPEC (0.50);Not determined (0.50) |
| 1187 | B1 | 15 | ST940 (1.00) | 0 | - | - | - |
| 125 | A | 13 | ST617 (0.92);ST4981 (0.08) | 7 | ST617 (1.00) | Americas (0.71);Europe (0.29) | ExPEC (0.57);Not determined (0.43) |
| 1188 | A | 12 | ST167 (1.00) | 0 | - | - | - |
| 1110 | F | 11 | ST1722 (1.00) | 1 | ST1722 (1.00) | Europe (1.00) | ExPEC (1.00) |
| 288 | A | 9 | ST5824 (1.00) | 3 | ST227 (1.00) | Americas (1.00) | Not determined (1.00) |
| 20 | B2 | 8 | ST131 (1.00) | 48 | ST131 (0.96);ST5432 (0.02);ST5494 (0.02) | Europe (0.75);Americas (0.12);Unknown (0.10);Oceania (0.02) | ExPEC (0.71);Not determined (0.29) |
| 37 | D | 8 | ST405 (1.00) | 27 | ST405 (0.96);ST964 (0.04) | Europe (0.63);Americas (0.33);Asia (0.04) | ExPEC (0.81);Not determined (0.11);ETEC (0.04);STEC (0.04) |
| 1189 | A | 8 | ST1421 (1.00) | 0 | - | - | - |
| 7 | D | 7 | ST69 (1.00) | 174 | ST69 (0.94);ST106 (0.03) | Europe (0.83);Americas (0.12);Unknown (0.05) | ExPEC (0.79);Not determined (0.19);EAEC (0.02) |
| 1190 | A | 7 | ST9847 (1.00) | 0 | - | - | - |
| 1191 | D | 6 | ST38 (0.83);Novel (0.17) | 0 | - | - | - |
| 92 | A | 5 | ST10 (1.00) | 10 | ST10 (1.00) | Europe (0.60);Americas (0.40) | ExPEC (0.60);Not determined (0.40) |
| 164 | A | 5 | ST34 (1.00) | 6 | ST34 (1.00) | Africa (0.67);Europe (0.33) | EPEC (0.67);Not determined (0.33) |
| 330 | B1 | 5 | ST101 (1.00) | 2 | ST101 (1.00) | Europe (0.50);Unknown (0.50) | Not determined (1.00) |
| 1193 | A | 5 | ST10 (1.00) | 0 | - | - | - |

**Figure 2:**
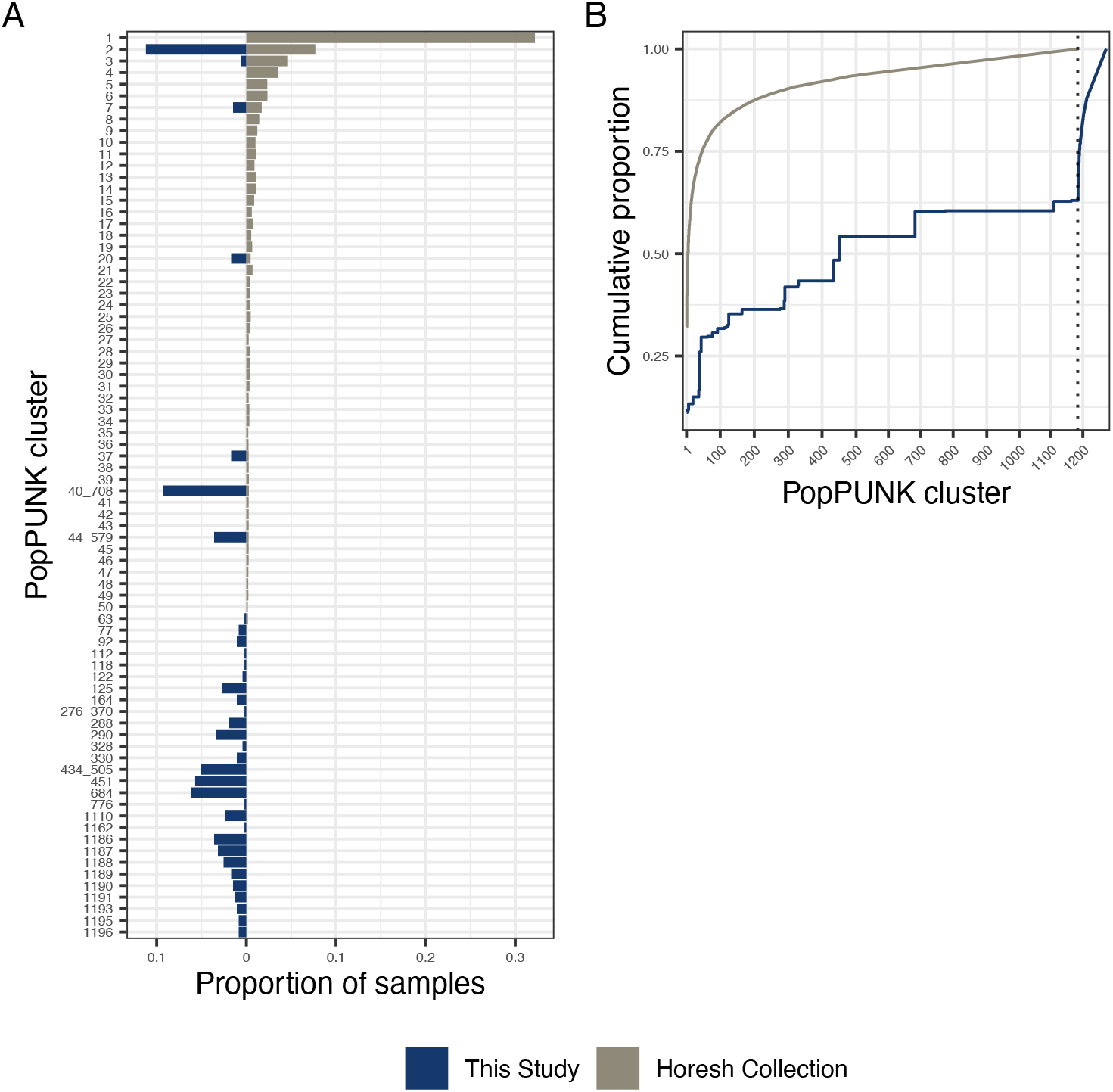
Comparing distribution of popPUNK clusters in Malawian and global collections. A: proportion of samples assigned to a given popPUNK cluster in Malawian (left) and global (right) isolates. Clusters are arranged in name-numeric order which, by definition, is size order from the original publication from largest to smallest. Clusters 1-50 (accounting for 76% of global isolates) are shown along with any cluster containing at least 3 Malawian isolates. B: Cumulative proportion of samples with given cluster membership, stratified by study; clusters are again numerically ordered on x-axis. Dotted line shows the maximum cluster identifier that was defined in the global collection (n = 1184); clusters with an identifier greater than 1184 (to right of dotted line) were not present in global collection. Clusters with an identifier made up of two numbers separated by an underscore are clusters that were two separate clusters in the original global collection but have been merged after Malawian genomes were added (e.g. 40_708).

We next reconstructed a core-gene phylogeny using all 473 genomes from our collected isolates, 2,776 assemblies from the global collection (selected to span the diversity of the collection) and 97 genomes from a previous study at QECH representing mostly invasive isolates from the same setting as our carriage collection. (Figure 3). Malawian isolates were distributed throughout the tree, suggesting that Malawian strains represent a subsample of the global diversity of *E. coli*. We observed apparent clonal expansions of Malawian isolates in ST410, ST167 and ST131 (Figures 3B-D), though in the case of ST131 (Figure 3D) this included one isolate from the USA (year unknown).

**Figure 3:**
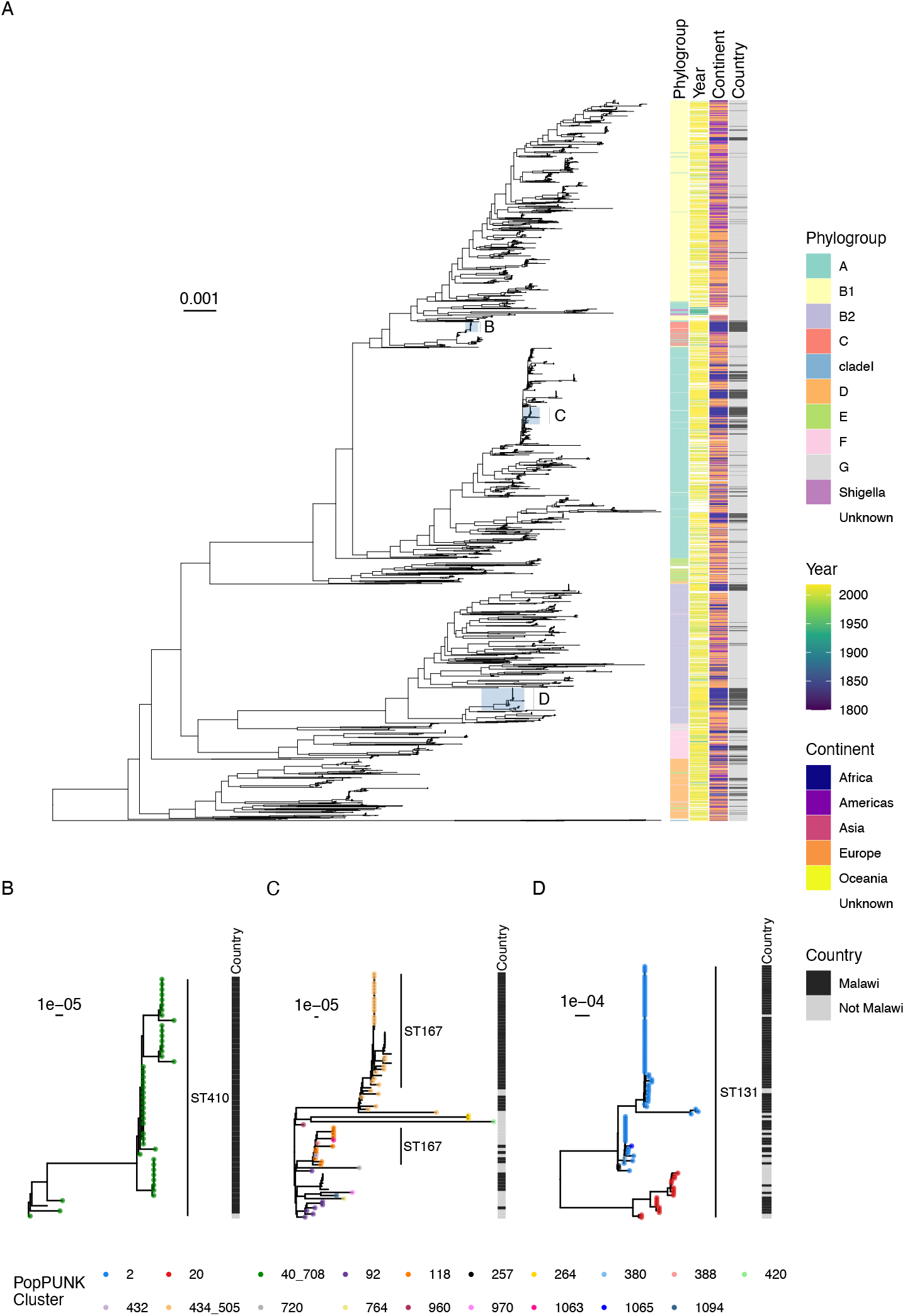
Midpoint rooted core-gene maximum likelihood phylogenetic tree of Malawian isolates with global context isolates (10 isolates each from the top 50 popPUNK clusters in the global collection, along with a further 92 Malawian isolates from a previous study), showing phylogroup, year of collection, continent, and country (Malawi vs not Malawi). Generally, Malawian isolates are distributed throughout the tree. B-D show magnified subtrees of the three most frequently identified STs: ST 410 (B), ST167 with surrounding phylogroup A isolates (C) and ST131 (D) with tip points coloured based on popPUNK cluster allocation. Lack of coloured point indicates that the isolate was assigned a novel cluster not present in the original global collection.

To explore the genomic epidemiology of ST131, ST410 and ST167 with higher resolution, we reconstructed phylogenies by mapping each ST dataset to ST-specific reference genomes, incorporating 140 ST167, 371 ST410 and 919 ST131 genomes (Supplementary Figures 3 and 4 and Figure 4), confirming our samples forming distinct subclades in ST167 and ST410. The Malawian ST410 (except for two isolates) are part of the globally distributed, often carbapenemase-associated B4/H24RxC lineage and formed a well-supported (>95% UltraFast bootstraps) monophyletic clade within the B4/H24RxC lineage (Figure 4A), with a median 35 (range 0-60) SNPs between the 43 Malawian isolates. All except one ST167 isolates comprised two lineages, one of which formed a well-supported (>95% UltraFast bootstraps) monophyletic clade of only Malawian isolates (Figure 4B) and a median 24 (range 0-66) SNPs between the 24 isolates; the other lineage included Malawian and global isolates. The Malawian ST131 isolates were, in contrast, distributed throughout the ST131 phylogeny (Figure 4C).

**Figure 4:**
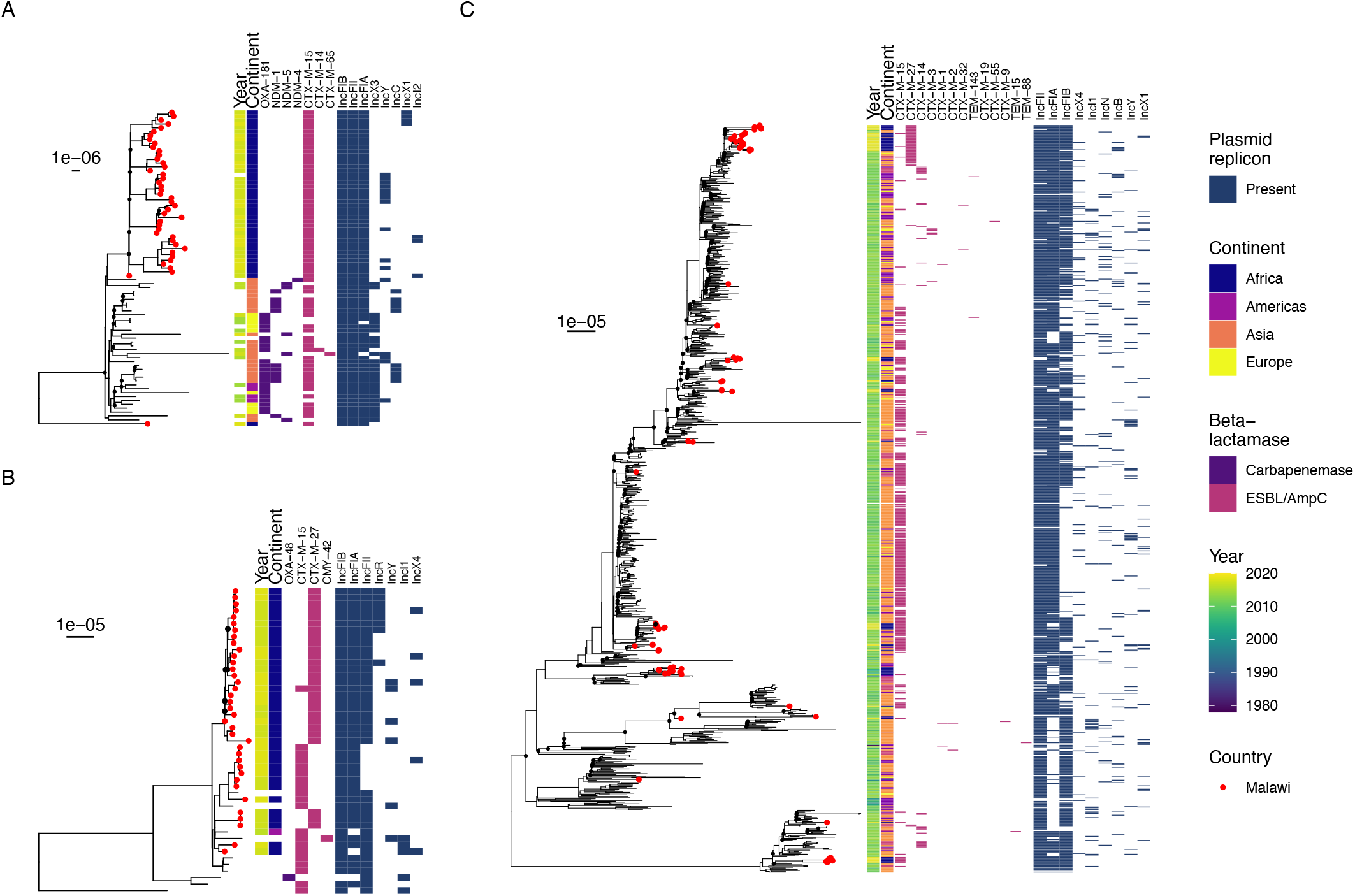
Subtrees of midpoint-rooted, maximum likelihood phylogenies of global *E. coli* ST410 (A) and ST167 (B) collections, and the global ST131 phylogeny (C) showing Malawian isolates (red tip points). Assemblies were constructed by mapping to ST-specific reference genomes. ESBL/CPE genes and plasmid replicons are shown. Bootstrap support of less than 95% is shown by a black point at tree node. Full ST 410 and ST167 global trees are shown in Supplementary Figures 3 and 4.

**Figure 5:**
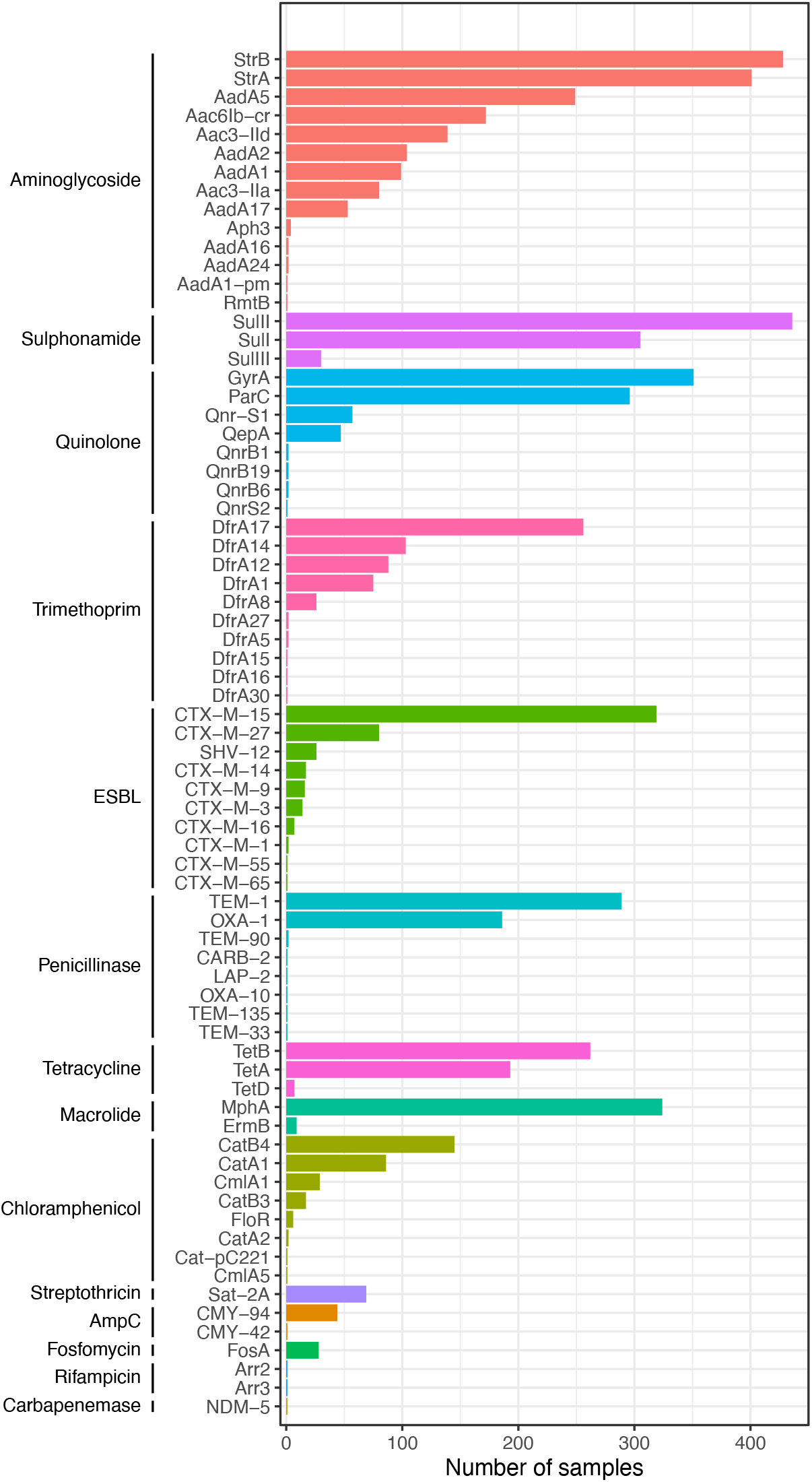
Distribution of identified antimicrobial resistance determinants

### Resistance and virulence determinants

We identified a variety of AMR determinants in the isolates sequenced for this study (Figure 5). One ST2083 isolate contained a carbapenemase encoding gene, *bla*_NDM-5_, which, in our short read *de novo* assembly, was co-located with an Inc-X3 containing contig, consistent with carriage on a plasmid. The ST410 B4/H24RxC lineage is often associated with a characteristic *bla*_NDM-5_-associated Inc X3 plasmid (pNDM5_020026)^8^ similar to our ST2083-IncX3-*bla*_NDM-5_ contig, as we described previously ^14^. To further characterise our ST2083 isolate we used long-read sequencing, to generate a hybrid assembly for this sample, as well as for a randomly selected ST410 isolate of the B4/H24RxC-like lineage. This confirmed that our ST410 isolate carried two of the four ST410 B4/H24RxC-associated plasmids, including the *bla*_CTX-M-15_ carrying pCTXM15_020026 plasmid, but lacks the *bla*_NDM-5_ and IncX-encoding pNDM5_020026 plasmid (Supplementary Figure 5). However, hybrid assembly of the sample from ST2083 showed that it carries a plasmid almost identical to the pNDM5_020026 (Supplementary Figure 5; using the default BRIG cutoff of 70% similarity to display a sequence section as conserved), highlighting the potential risk of this plasmid should it move into one of the more successful STs, in particular the ST410 B4/H24RxC lineage also commonly identified in our study (Supplementary Figure 5).

The remainder of the isolates without carabenemases (n = 472) carried at least one ESBL-encoding gene, most commonly *bla*_CTX-M-15_ which was present in 319/473 (67%) of isolates. All other identified ESBL-encoding genes were members of the *bla*_CTX-M_ family except for *bla*_SHV-12_ which was identified in 26/473 isolates across 6 STs, but was particularly common within ST656; all 17 ST656 isolates in the collection carried *bla*_SHV-12_ (Supplementary Figure 6)

Co-occurring determinants of resistance to aminoglycosides (99% [472/473] of isolates), trimethoprim (97% [459/473]), sulphonamides (99% [468/473]) and quinolones were very common (86% [407/473]), whilst genes conferring resistance to chloramphenicol were of lower frequency (52% [248/473]). The most frequently identified quinolone resistance determinants were QRDR mutations (*gyrA* in 74% [351/473] and *parC* in 63% [296/473] isolates), but plasmid-mediated quinolone resistance determinants were also found (*qnrS* [12%, 58/473], *qnrB* [1% 6/473] and *qep* [10%, 47/473]). The *gyrA* mutants were S83L (n=351), D87N (n= 294) and *parC* mutants S80I (n=296) and S84G (n=6); co-occurrence of QRDR mutations and *qnr* genes was unusual (6 isolates). Most (461/473 [97%]) strains lacked the genes to classify them as any pathotype: 2/473 were identified as aEPEC/EPEC and 10/473 as EAEC (Supplementary Figure 1).

## Discussion

Understanding how the genomic epidemiology of *E. coli* in multiple LMICs differs from high income settings is essential to provide insight into local and global drivers of transmission. Our genomic analysis of the diversity of ESBL-producing *E. coli* in Blantyre, Malawi, enhances our understanding of the global *E. coli* genomic epidemiology by adding data from an under-sampled location. ESBL-producing *E. coli* colonising adults in Malawi represents the diversity of the species with all major phylogroups, and 57 STs are represented. Some global trends in ST are broadly reflected in Blantyre; ST131, for example, the most frequently isolated pathogenic ST in many settings worldwide^47^, was the most commonly isolated ST, followed by the globally emergent AMR-associated ST410 and ST167^7,8^. By placing the Malawian isolates in a wider context we found diversity in the Malawian collection not captured in the curated species-wide Horesh collection^9^. There was a difference in relative prevalence of lineages as defined by popPUNK between Malawian and other samples, with many Malawian isolates forming distinct clusters. Both the core gene and map-to reference phylogenies (using different context genomes) were consistent with local Malawian subclades in ST410 and ST167. This could be due to true locally circulating subclades or sampling biases in global *E. coli* collections either spatially (i.e. sampling high income settings) or particular lineages (e.g. sampling ST131 or other important pathogens) or resistance phenotypes (e.g. we have selected for ESBL producers); only 246/10,146 genomes in the global *E. coli* dataset used to build the curated collection in Horesh et al. were from the African continent^9^.

Scaling up of genomic surveillance in sSA with sampling of human, environmental and animal isolates can redress this imbalance, improve understanding of the global transmission of AMR *E. coli*, and should be an international priority for funders of future studies. Efforts to expand global collections (such as the species-wide collection used here) in an unbiased manner as further genomes from low- and middle-income countries are sequenced will also be key to developing a comprehensive picture of local, national, and global scales of AMR transmission.

In Blantyre, as worldwide, *bla*_CTX-M_ is the dominant ESBL family, especially *bla*_CTX-M-15_. In our isolates, which were selected based on ESBL production, only one isolate carried a carbapenemase encoding gene: *bla*_NDM-5_. As observed commonly in collections of ESBL-producing bacteria dependent on mobile elements, aminoglycoside, trimethoprim and sulphonamide resistance determinants were near-universal, and ciprofloxacin and chloramphenicol resistance determinants common. In terms of plasmids the co-occurrence of *bla*_CTX-M-15_ and IncF plasmids in our isolates reflects predominant global observations.^48,49^

This AMR-determinant distribution may be influenced by local antimicrobial pressures: carbapenem antimicrobials are at best sporadically available in QECH, but co-trimoxazole as CPT is widely used in this high HIV-prevalence setting as lifelong prophylaxis against infection in people living with HIV, as per World Health Organisation guidelines^16^. The ubiquity of genes conferring resistance to co-trimoxazole in this collection (i.e. *dfr, sul*) raises the possibility that use of CPT is creating selection pressure for other AMR-determinants in Blantyre. CPT has been shown to reduce mortality due to bacterial infection and malaria^16^ in people living with HIV in high-burden settings. However, in an era of increasingly effective HIV treatment and increasing Gram-negative resistance it may be that a more nuanced approach to the deployment of CPT is needed.

ESBL producing Gram-negative infections are an increasing clinical problem in Malawi, and there is a significant unmet need for access to carbapenem antimicrobials to treat them. Carbapenemase producing Enterobacterales including *E. coli* (*bla*_NDM-5_ in *E. coli* ST636) have recently been described in other regions of Malawi^50^, and *bla*_NDM-5_ is increasingly being identified across sSA^51^. The presence of carbapenemases in this collection despite minimal local carbapenem use highlights the need for ongoing surveillance as these last-line antimicrobials are introduced. Globally, ST167 and ST410 include carbapenemase-associated lineages, but carbapenem-associated ST410 and 167 have not yet been observed in our setting. Amongst ST410, our isolates are however closely related to the globally distributed B4/H24RxC *bla*_NDM-5_/*bla*_OXA-181_ carbapenemase-associated lineage but lacked carbapenemase genes. Long read sequencing demonstrates however that the characteristic B4/H24RxC *bla*_NDM-5_-associated plasmid is almost identical to the plasmid observed in our collection in a different sequence type (ST2083), highlighting how important it is to introduce surveillance of emerging carbapenemase-expressing bacteria. Lack of carbapenemase encoding genes in our B4/H24RxC isolates could be due to lack of acquisition of harbouring plasmids, or due to repeated acquisition and subsequent loss events given the current absence of carbapenem selection pressure. If the latter it would suggest the fitness trade-off to harbour the plasmid might select against maintenance in the absence of the selection pressure within this lineage for now. Availability of carbapenems is predicted to change.

Chloramphenicol has historically been a first-line treatment for severe febrile illness in Malawi^52^ but increasing numbers of resistant isolates causing infections has curtailed its use in favour of ceftriaxone^53^. Chloramphenicol resistance determinants were however absent in 48% of our samples; indicating that chloramphenicol could still have a role to play as a reserve agent in the treatment of ESBL-*E. coli*, but this approach would require quality assured diagnostic microbiology services to support it.

There are limitations to our study. Some participants provided multiple samples so it is possible that this introduced bias into the collection. Our study was predominantly based at a single urban hospital, over a time period of around only 2 years so although it contributes valuable genomes from an under sampled region and population, the extent to which these genomes are representative of other areas on Malawi is unclear. Furthermore, our samples were cultured on ESBL-selective media, meaning we provide a snapshot of ESBL positive *E. coli* genomic diversity in Malawi, rather than of *E. coli* infecting humans as a whole, which contrasts with the Horesh study, which did not impose the same selection criteria.

In conclusion, we find that the diversity of ESBL *E. coli* from Blantyre includes globally distributed high-risk clones and broadly reflects the global population structure. The occurrence of distinct monophyletic subclades when assessed in the context of a large recent data collection highlights the need for further targeted sequencing of isolates from Malawi and sSA to understand local, regional, and global *E. coli* circulation. Carbapenemase-associated lineages reported elsewhere in the world are present in Malawi, but currently still lack carbapenem resistance determinants. However, the presence of carbapenemase-encoding plasmids in our collection highlights that it is likely only a matter of time before either these lineages are selected for and evolve into high-risk clones or carbapenemase genes on mobile genetic elements are transmitted to a high-risk clone. There is a critical need for both robust stewardship strategies and for ongoing genomic surveillance with rapid reporting, as these agents are introduced.

## Supporting information

Sample accession and metadata

Metadata for Horesh global context isolates

Accessions and metadata for ST131 context isolates

Accessions and metadata for ST167 context isolates

Accessions and metadata for ST410 context isolates

## Author contributions

Conceptualisation: JML, NT, NF, JM; Data curation: JL; Formal analysis: JL, MAB, AR, EH, NRT, NAF; Funding acquisition: JML, NRT, NAF; Investigation: JML, MM, RB, CA, AZ, EH; Resources: NRT; Writing – original draft: JL; Writing – review and editing: all authors

## Ethical approval

The clinical study which provided the isolates for this analysis was approved by the Liverpool School of Tropical Medicine (16-062) and Malawi College of Medicine (P.11/16/2063) research ethics committees.

## Acknowledgments

Many thanks to the study nurses Lucy Keyala, Tusekile Phiri, Grace Mwaminawa, Witness Mtambo, Gladys Namacha, Monica Matola; to the MLW laboratory teams, particularly Brigitte Denis; and to the MLW data team, particularly Lumbani Makhaza and Clemens Masesa. The authors acknowledge the sequencing team at the Wellcome Sanger Institute, and Christoph Puethe and the Pathogen Informatics team for computational support. This research was funded in whole, or in part, by the Wellcome Trust (#206194 and 108413/A/15/D). For the purpose of open access, the authors have applied a CC-BY public copyright licence to any author-accepted manuscript version arising from this submission.

