## Supplementary Figures and Tables for "Genomic analysis of extended-spectrum beta-lactamase (ESBL) producing *Escherichia coli* colonising adults in Blantyre, Malawi reveals previously undescribed diversity"

### Supplementary material

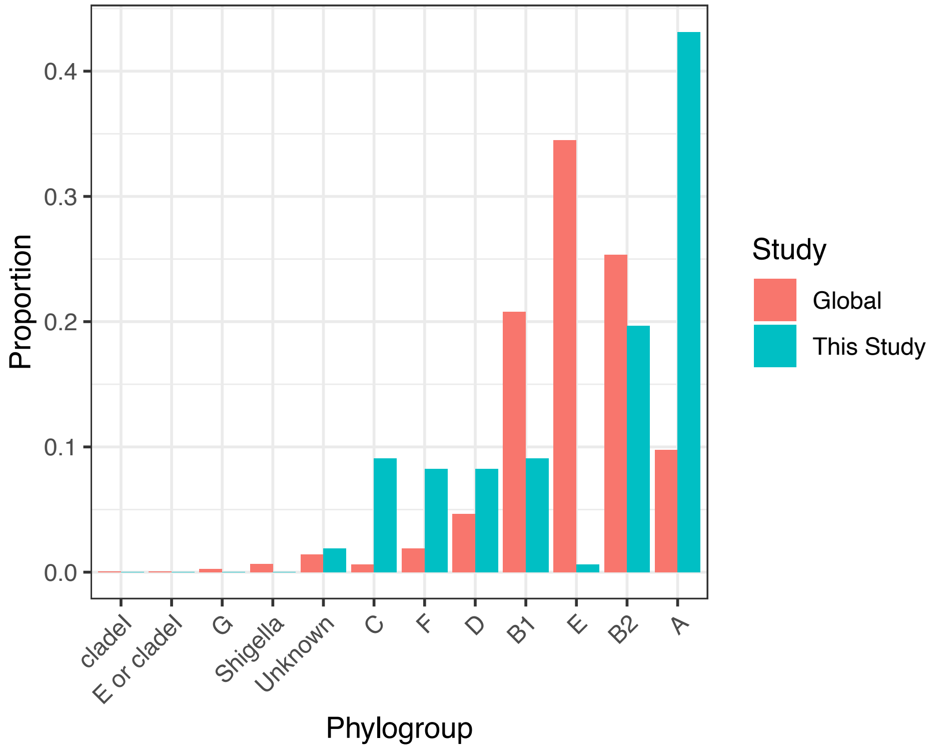


**Supplementary Figure 1:** Comparison of phylogroup distribution between this study and the global collection of *E. coli* isolates.

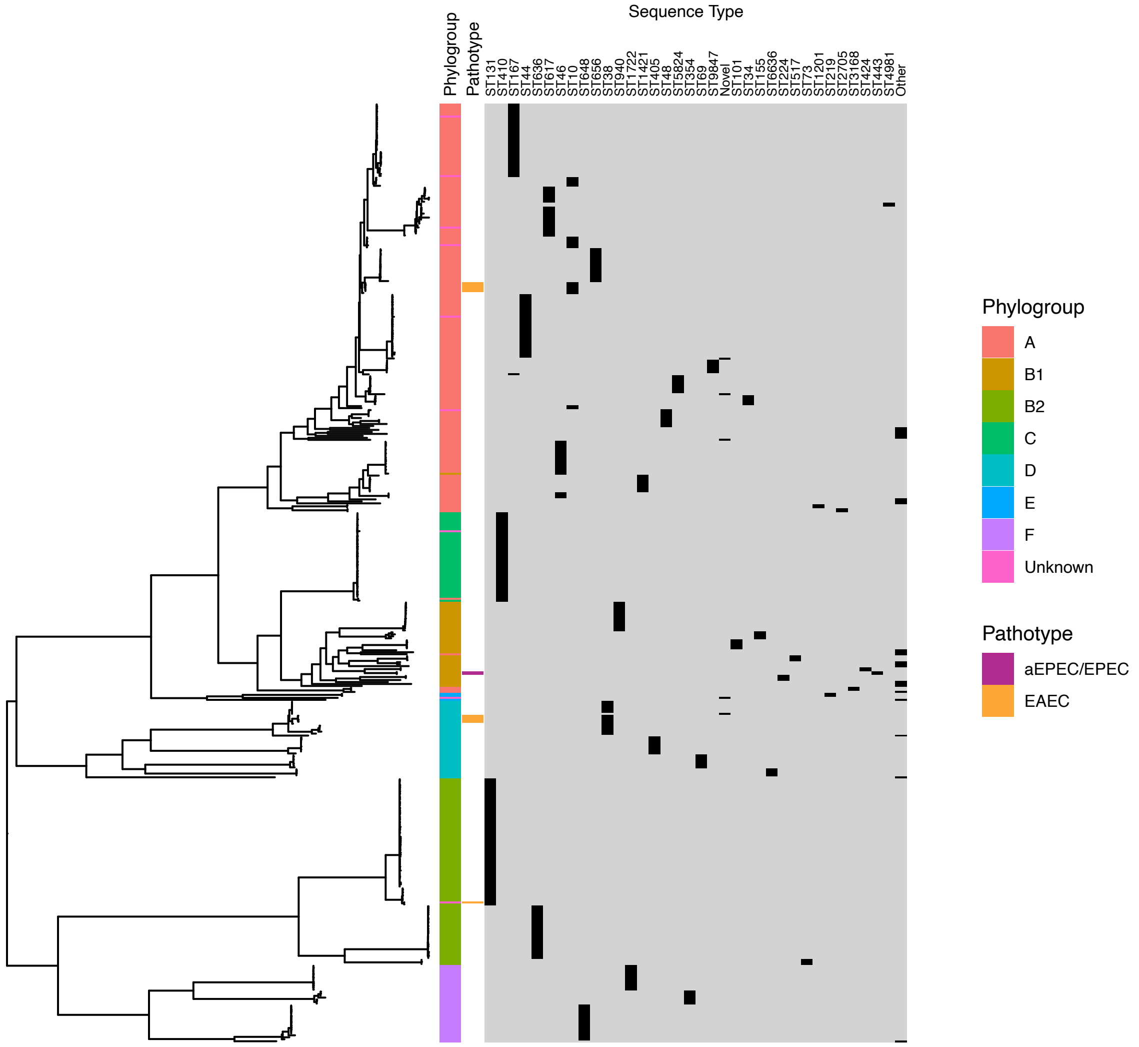


**Supplementary Figure 2:** Phylogeny of E. coli from this study. Maximum-likelihood midpoint-rooted phylogeny inferred using the general time-reversible nucleotide substitution model with FreeRate site heterogeneity with five parameters, as selected by IQTREE ModelFinder module. Phylogroups, STs and pathotypes shown.

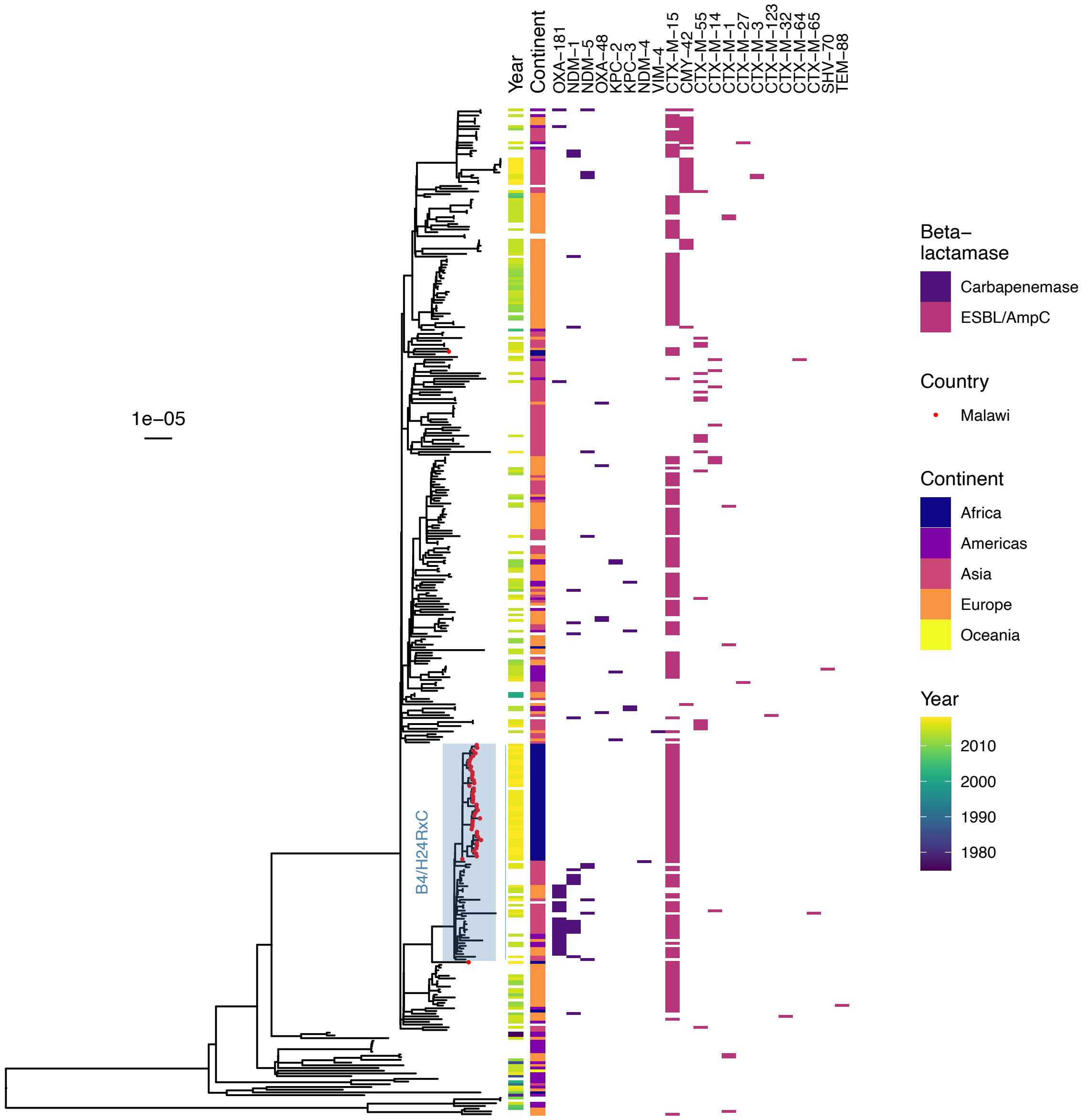


**Supplementary Figure 3:** Midpoint-rooted maximum likelihood phylogeny of global E. coli ST410, with assemblies obtained by mapping to reference. ESBL/CPE genes and plasmid replicons are shown. Blue shaded area shows the carbapenemase-associated B4/H24RxC lineage – this area is expanded in Figure 4.

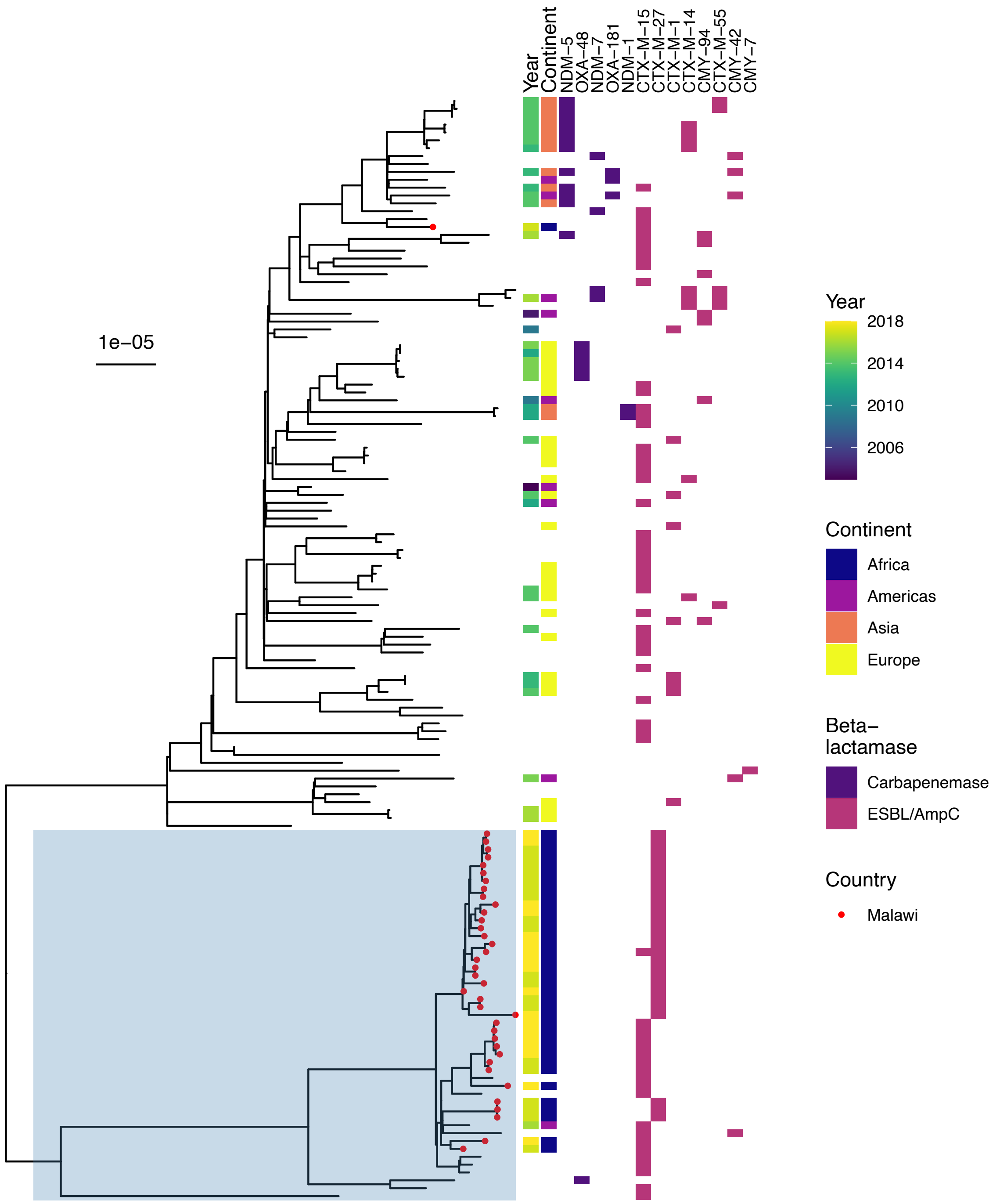


**Supplementary Figure 4:** Midpoint-rooted maximum likelihood phylogeny of global *E. coli* ST167, with assemblies obtained by mapping to reference. ESBL/CPE genes and plasmid replicons are shown. Blue shaded area shows the area that is expanded in the subtrees in Figure 4.

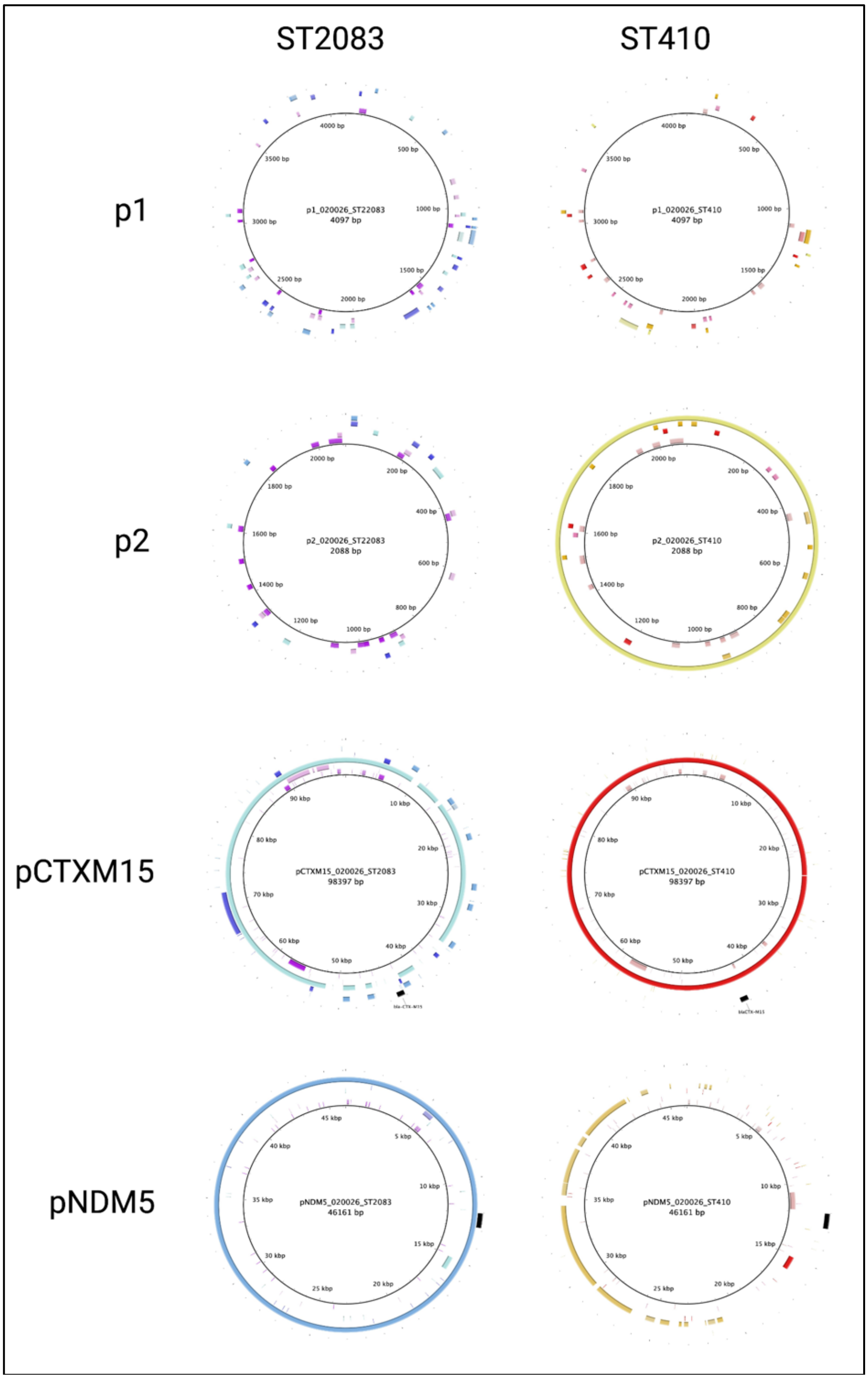


**Supplementary Figure 5:** Blast Ring Image Generator (BRIG) comparison of the plasmids present in the ST410 B4/H24RxC MDR isolate 020026 from Feng at al[8] with the ST2083 (left panels) and ST410 (right panels) assemblies from our study. Inner to outer ring show the different plasmids from the respective isolate (ST2083 or ST410). Whilst ST410 seems to share the identical plasmid p2 (right panel second row) and the blaCTX-M-15 encoding plasmid (right panel third row), the bla-NDM-5 encoding plasmid is only partially conserved; of particular importance, the bla-NDM-5 gene (indicated as black box outside all rings) is not present, as predicted by ariba. On the contrary, the ST22083 isolate has only partial similarity with the bla-CTX-M-15 encoding plasmid (third row, left column) but the completely conserved bla-NDM-5 encoding plasmid, including the NDM-5 gene (left panel, row four).

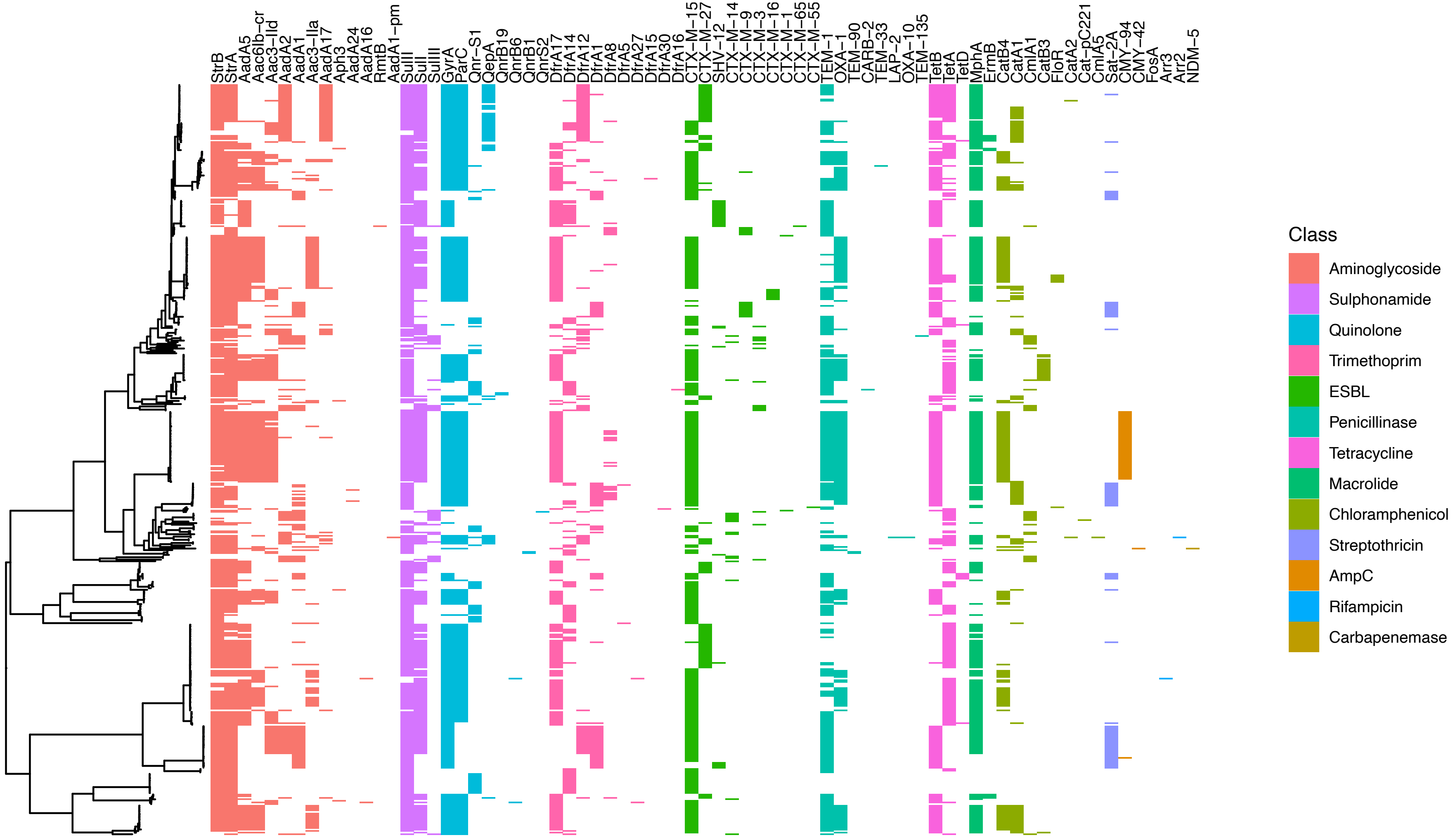


**Supplementary Figure 6:** Antimicrobial resistance determinants mapped back to core gene tree

**Supplementary Table 1:** *E. coli* pathotype definitions

| **Definition** | **Pathotype** |
|---|---|
| Presence of any Shiga toxin gene | STEC (Shiga toxin producing *E. coli*) |
| Presence of *eae* | aEPEC/EPEC ([atypical] Enteropathogenic *E. coli*) |
| Presence of Shiga toxin plus *eae* | EHEC (Enterohaemorrhagic *E.coli*) |
| Presence of *aatA or aggR or aaiC* | EAEC (Enteroaggregative *E. coli*) |
| Presence of *est* or *elt* | ETEC (Enterotoxigenic *E. coli*) |
| Presence of pINV plasmid | EIEC (Entero invasive *E. coli*) |
